# Collective polarization dynamics in gonococcal colonies

**DOI:** 10.1101/2022.08.04.502895

**Authors:** Marc Hennes, Niklas Bender, Tom Cronenberg, Anton Welker, Berenike Maier

**Affiliations:** Institute for Biological Physics, and Center for Molecular Medicine Cologne, University of Cologne, Cologne, Germany

**Keywords:** membrane potential, biofilm, bacterial colony, antibiotic tolerance, collective dynamics

## Abstract

Membrane potential in bacterial systems has been shown to be dynamic and tightly related to survivability at the single cell level. However, little is known about spatio-temporal patterns of membrane potential in bacterial colonies and biofilms. Here, we discovered a switch from uncorrelated to collective dynamics within colonies formed by the human pathogen *Neisseria gonorrhoeae*. In freshly assembled colonies, polarization is heterogeneous with instances of transient and uncorrelated hyper- or depolarization of individual cells. As colonies reach a critical size, the polarization behaviour switches to collective dynamics: A hyperpolarized shell forms at the centre, travels radially outward, and halts several micrometres from the colony periphery. Once the shell has passed, we detect an influx of potassium correlated with depolarisation. Transient hyperpolarization also demarks the transition from volume to surface growth. By combining simulations and the use of an alternative electron acceptor for the respiratory chain, we provide strong evidence that local oxygen gradients shape the collective polarization dynamics. Finally, we show that within the hyperpolarized shell, tolerance against aminoglycoside antibiotics but not against β-lactam antibiotics is increased, suggesting that depolarization instantaneously protects cells, while the protective effect of growth arrest does not set in immediately. These findings highlight that the polarization pattern can demark the differentiation into distinct subpopulations with different growth rates and antibiotic tolerance.

**Significance statement:** At the level of single bacteria, membrane potential is surprisingly dynamic and transient hyperpolarization has been associated with increased death rate. Yet, little is known about the spatiotemporal dynamics of membrane polarization during biofilm development. Here, we reveal a discrete transition from uncorrelated to collective polarization dynamics within spherical colonies. Suddenly, a shell of hyperpolarized cells forms at the colony centre and hyperpolarization travels radially outward. In the wake of this shell, bacteria depolarize, reduce their growth rate, and become tolerant against antibiotics, indicating the onset of habitat diversity. Single cell live imaging and modelling link hyperpolarization to an oxygen gradient formed within the colonies. We anticipate that dynamical polarization patterns are tightly connected to biofilm differentiation in various bacterial species.

## Introduction

Bacteria actively maintain a negative membrane potential as part of the ion motive force. Functioning as a source of energy, ion motive force powers ATP synthesis, transport across the membrane, and membrane-standing molecular machines (1-3). Recent studies investigating membrane potential of *Escherichia coli* and *Bacillus subtilis* at the single cell level revealed that the membrane potential is highly dynamic and heterogeneous (4, 5). Transient hyperpolarization is associated with increased death rate (6), while depolarization has been shown to maintain viability under oxygen depletion (7) and antibiotic treatment. In particular, aminoglycoside antibiotics induce hyperpolarization (6, 8) and hyperpolarized cells tend to grow more slowly and have a higher death rate (6). These reports provide evidence that inhibition of hyperpolarization maintains growth and viability. While there is no evidence of collective hyperpolarization in bacterial populations, spatially propagating waves of membrane depolarization have been found in *B. subtilis* colonies (9). These waves coordinate the metabolic state and growth behaviour between the interior and the periphery through the release of intracellular potassium via dedicated ion channels (9, 10). To our knowledge, collective dynamics of membrane potential in colonies and biofilms has not been found in other species so far.

Biofilms are an abundant form of bacterial life (11). Within biofilms, localized gradients of nutrients, oxygen, and waste provide habitat diversity (12). One consequence of this diversity is that bacteria partition into fast growing cells at the surface of the biofilm and slowly growing cells at the centre (12, 13). Slow growth tends to increase tolerance of the bacteria against different antibiotics and other stresses (12, 14). Heterogeneity and dynamics of membrane potential potentially contribute to local habitat formation and antibiotic tolerance. Well studied biofilm-formers like *Pseudomonas aeruginosa* initiate biofilm formation by surface attachment of single planktonic cells and subsequent proliferation into colonies (15, 16). The irreversible switch from the planktonic state into the biofilm state is believed to occur gradually at this stage (16). In this study, we address these dynamics using *Neisseria gonorrhoeae* (gonococcus), the causative agent of the second most prominent sexually transmitted disease, gonorrhea (17). By contrast to *P. aeruginosa, N. gonorrhoeae* uses type 4 pilus (T4P) driven motility to self-aggregate into surface-attached spherical microcolonies consisting of thousands of cells within a few minutes (18-20). T4P are extracellular polymers that continuously elongate and retract (21-24). T4P dynamics are crucial for the structure of gonococcal colonies; a tug-of-war mechanism fluidizes the colonies, introducing local liquid-like order and causing colonies to form spheres (25-30). Colony formation protects *N. gonorrhoeae* against the β-lactam ceftriaxone (31) and the degree of tolerance depends on the physical properties of the colonies (29). Within several hours, a gradient of growth rates develops in these colonies (32), but it is unclear how and at which time scale habitat diversity emerges. Such transitions are expected to occur at some point in the maturation process of the biofilm, and may be linked to cell differentiation and the emergence of sleeper cells (12, 14). The spherical geometry makes gonococcal colonies an ideal system for studying the evolution of the membrane potential during maturation of freshly assembled colonies into biofilms.

In this study, we focus on the dynamics of membrane potential in gonococcal colonies and reveal a switch from a loner state to a collective state. Using single cell analysis within spherical colonies, we investigate the polarization dynamics at different stages of colony development. In freshly assembled colonies, single cell membrane potential is heterogeneous and spatially uncorrelated. Eventually, a shell of hyperpolarized cells occurs at the colony centre and travels towards the colony periphery. This event signifies a switch to collective membrane potential dynamics and correlates with reduction of growth. Behind the hyperpolarized shell, the intracellular potassium concentration increases and cells depolarize. A reaction-diffusion model strongly suggests that the dynamical pattern of oxygen concentration shapes the polarization dynamics. Application of the protein synthesis inhibitors kanamycin or azithromycin reverses the direction of the traveling shell. Within the shell, tolerance against kanamycin increases. Taken together, we show that membrane potential dynamics signifies a switch to collective behaviour in gonococcal colonies.

### In freshly assembled colonies, the membrane potential of neighbouring cells is uncorrelated

Driven by type 4 pili, *N. gonorrhoeae* cells actively assemble into colonies comprising thousands of bacteria (18, 29). We characterized the membrane potential of single cells within freshly assembled colonies growing in flow chambers that continuously provide fresh medium. The fluorescence intensity of the Nernstian dye TMRM was determined using confocal microscopy. Depending on the electrical potential *V*_*m*_ across the inner membrane, the cationic dye partitions between the cytoplasm and the extracellular space (4). The fluorescence intensities inside and outside of the cells were used to measure the membrane potential as described in the Methods. Since the membrane potential is negative, an increase in fluorescence intensity indicates an increase in polarization, i.e. the potential becomes more negative.

We found that the TMRM signal in young colonies is highly heterogeneous (Fig. 1a, Fig. S1). The derived membrane potential *V*_*m*_ of single cells shows a Gaussian distribution (Fig. 1b). Its mean value of *V*_*m*_ = −95 *mV* in slighly lower than the value obtained for single cells (33). With time, the mean membrane potential of cells in the colony decreases by roughly 15 mV, indicating steady depolarization of the population (Fig. 1b). The distribution of *V*_*m*_ departs from the initial Gaussian shape towards a distribution with heavier tails at larger potentials, attributable to a randomly distributed subpopulation maintaining strong polarization (Fig. 1a, b).

**Fig. 1.**
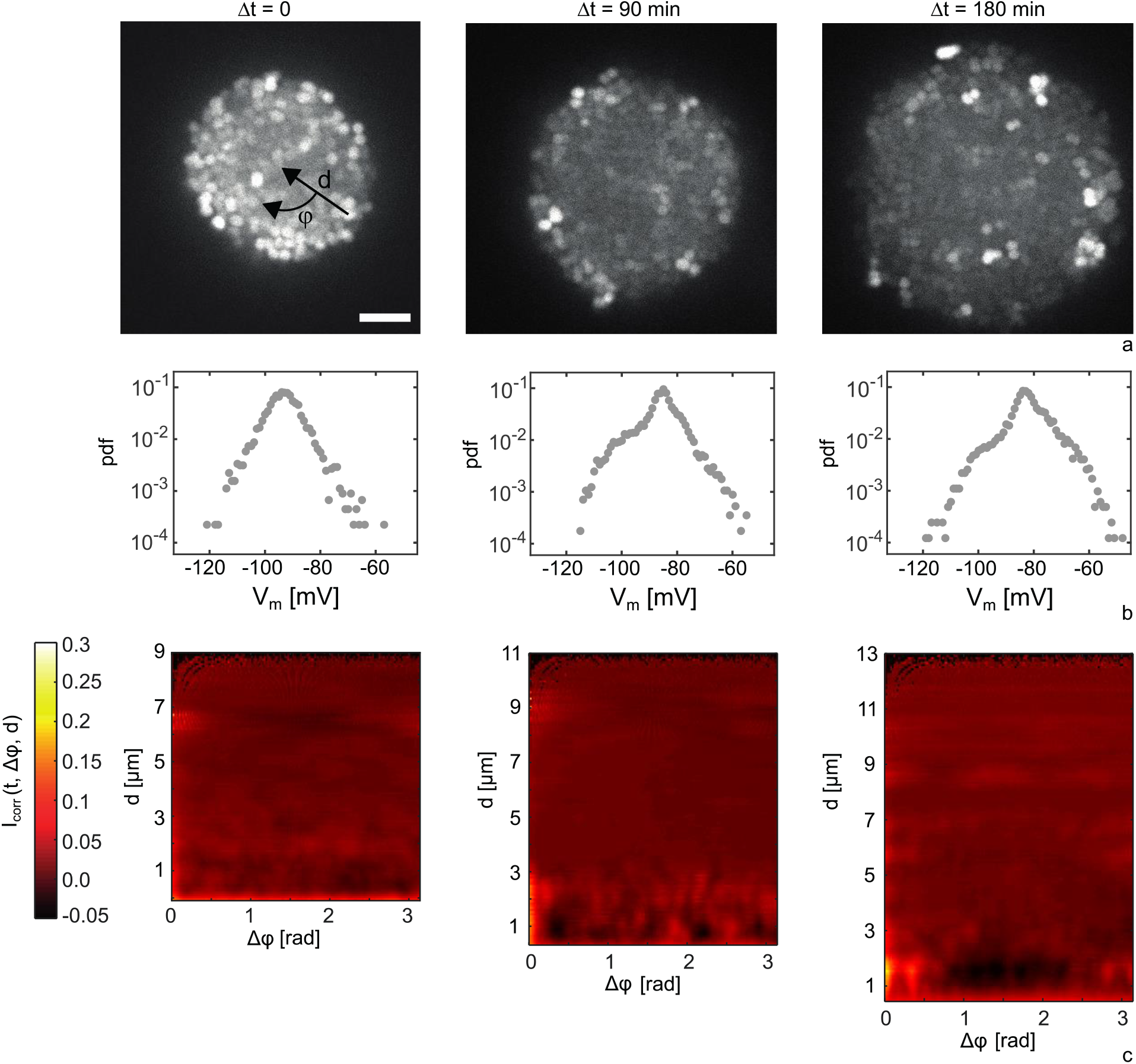
Heterogeneity of membrane potential in small colonies formed at different time points. Strain *wt green* (NG194), flow chamber. a) Typical fluorescence intensities from confocal slices through colony centre. Scale bar: 5 μm. b) Probability density function (pdf) of membrane potential of single cells at Δt = 0, 90 min, 180 min. (> 4000 cells for each time point). c) Angular correlation of the intensity fluctuations within the colony shown in a) 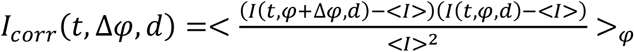 (colour coded) as a function of the angular position difference Δφ and the distance *d* from the edge of the colony.

At the single cell level, transient changes in polarization have been reported (4-6, 8, 34, 35). To find out whether the membrane potential is dynamic at the level of individual cells within colonies, we tracked the TMRM signal of single cells. To ensure that fluctuations in TMRM intensity are not caused by vertical movement of bacteria, we simultaneously tracked the sfGFP signal of the respective cells (strain *wt green*, see Table S1). A small fraction of cells experiences events of transient depolarization (Fig. S2a, b) or hyperpolarization (Fig. S2c, d, Movie S1) that last for ∼ 1-2 minutes. For all observed events, we find that transient changes in polarization (roughly 15 to 20 mV change) are uncorrelated between neighbouring cells. To quantify correlations in changes of the membrane potential, we defined a measure of how strongly deviations from the mean TMRM fluorescence intensity, (*t, x, y*) − ⟨*I*⟩, are spatially correlated between cells. ⟨*I*⟩ is the mean intensity of the colony at time t. In particular, we calculated the two-dimensional polar correlation function *I*_*corr*_(*t*, Δφ, *d*) = (*I*(*t, φ* + Δ*φ, d*) − ⟨*I*⟩)(*I*(*t, φ, d*) − ⟨*I*⟩)/⟨*I*⟩^2^ at different depths *d* of the colony (*d* = 0 μm at the edge). This function measures the correlation strength of the TMRM fluorescence signal in angular direction. We analysed *I*_*corr*_ for 20 colonies and found little correlation (Fig. 1c, further example in Fig. S3). This analysis confirms that the dynamics of transient changes in cellular membrane potential occurs independently between neighbouring cells.

We conclude that in freshly assembled colonies, the membrane potential of single cells is heterogeneous and spatially uncorrelated. Individual cells transiently hyper- or depolarize at the time scale of minutes while most of the population gradually depolarizes at a time scale of hours.

### In large colonies, a shell of hyperpolarized cells travels through the colony and marks the transition from volume to surface growth

As the colonies grew in size, we found that the membrane potential switched from the uncorrelated dynamics described above to collective dynamics. Sometime after colony assembly in flow chambers, cells in the centre of the colony showed increased polarization relative to the mean of the colony (Fig. 2a, Movie S2). Henceforth, we will refer to this relative increase in polarization as (collective) hyperpolarization. Collective hyperpolarization was transient and propagated radially from the centre towards the edge of the colony across all cells. Notably, the shell of hyperpolarized cells rarely reached the edge and mostly became stationary a few μm away from the interface. The angular correlation coefficient shows a clear maximum at the position of the hyperpolarized shell (Fig. 2b), indicating collective changes in membrane potential. The shell has a thickness of 3 - 4 μm, corresponding to 3 to 4 cell diameters. The difference of the membrane potential between the hyperpolarized cells and the cells at the colony centre is Δ*V*_*m*_ ≈ −10*mV* (Fig. 2c).

**Fig. 2.**
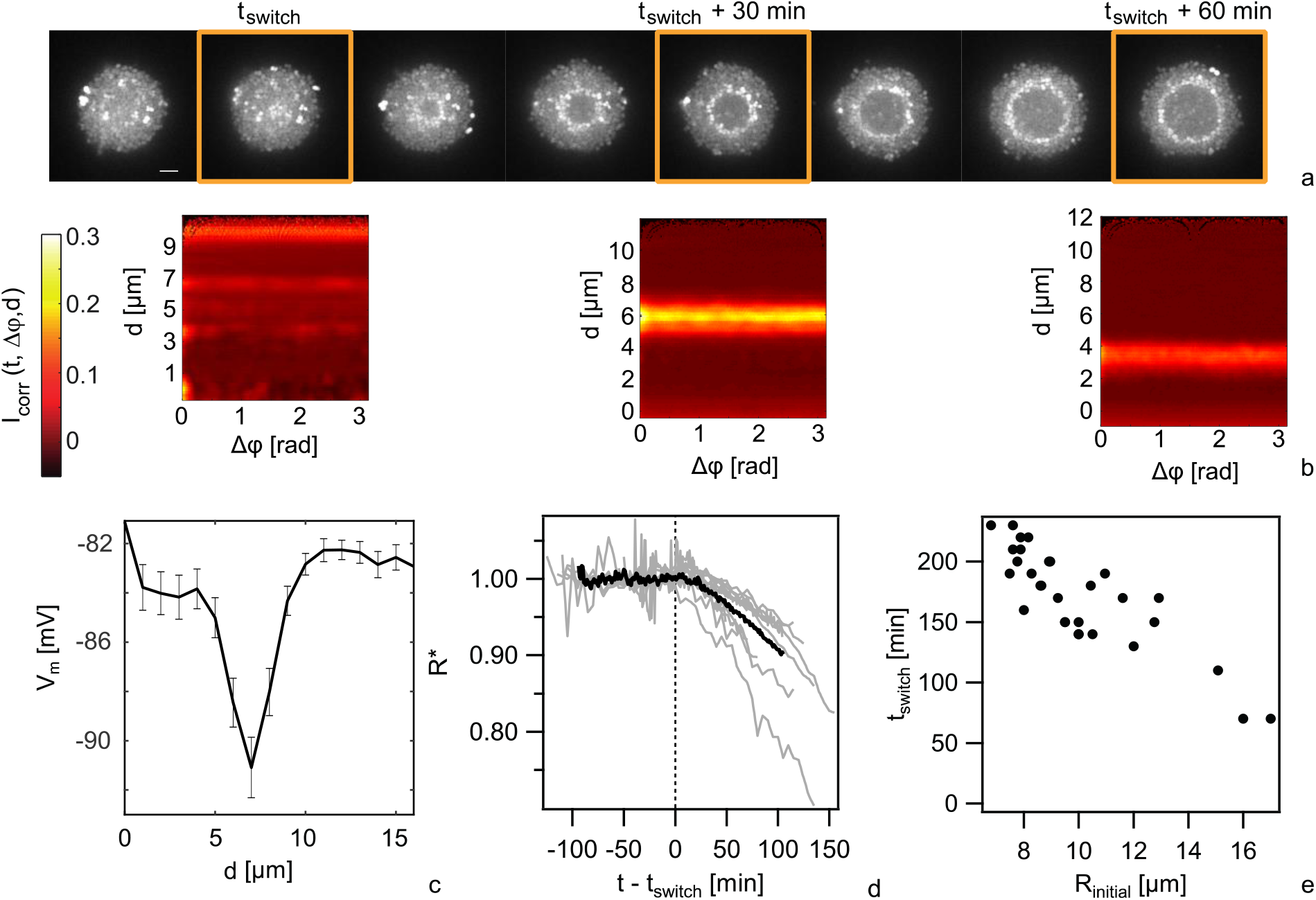
Travelling shell of hyperpolarized cells in large colonies. Strain *wt green* strain (NG194). *t*_*switch*_ is the time point at which the membrane potential switches to collective behaviour. a) Typical time lapse of TMRM fluorescence in flow cell, beginning 3 hours after incubation Scale bar: 5 μm. b) Angular correlation of the intensity fluctuations as a function of radial position within the colony 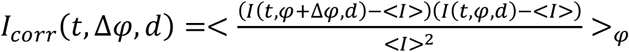. The images correspond to the fluorescence images in a) at the time points indicated by orange frames. c) Radial membrane potential of cells in colonies at the time point where the hyperpolarized cells reside 7 μm from the edge of the colony (*d*: distance from the edge of the colony). d) Colony radius *R*(*t*) normalized to an exponential function 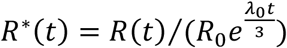. The time axis was shifted to set t = 0 at the time when collective hyperpolarization occurs in the TMRM image. λ_0_: initial growth rate, *R*_0_: initial colony radius. Deviation from 1 indicates deviation from exponential growth. Black line: Mean of 10 colonies from three different measurement days. Grey lines: individual colonies. e) Time point of switch to collective behaviour as a function of initial colony size in static cultures.

Previously, we showed that within colonies of ∼ 2 - 3 h of age, a characteristic pattern of growth rates forms whereby the rates decrease from the edge of the colony towards its centre (32). We assessed whether growth-inhibition and the onset of collective membrane dynamics are correlated. Deviation from an initially exponential growth would indicate reduced growth rates within colonies. We determined the radii of individual colonies as a function of time (start radius *R*_0_ and growth rate λ_0_) and normalized it by the radial growth function of the first 2 hours, yielding 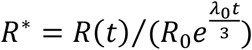. For the individual colonies, we found that the deviation from exponential growth coincided with the onset of collective hyperpolarization (Fig. 2d). Prior to the formation of the hyperpolarization shell (*t* − *t*_*switch*_ < 0), colony radii grew exponentially, *R*^*^ ≈ 1, with an average growth rate λ_0_ ≈ 0.6 *h*^−1^ in line with previous measurements (32). After shell formation (*t* − *t*_*switch*_ > 0), the normalized radius *R*^*^ quickly fell below 1. Relating the instantaneous radial position of the shell with the colony radius *R*(*t*), indicates that growth cessation is limited to regions of the colony through which the hyperpolarized shell has already passed (Supplementary Information, and Fig. S4). Previously, we showed that the growth rate is slowest at the colony centre while a narrow layer of cells continues to grow unperturbed (32). The layer of growing cells (32) and the layer of cells that has not experienced transient hyperpolarization both comprise 3 - 4 cell diameters. We conclude that collective hyperpolarization in our gonococcal colonies precedes growth arrest as observed at the single cell level (6), and marks the transition from volume to surface growth.

Within a single field of view in the microscope, the colonies do not switch simultaneously into the collective state (Movie S3). Larger colonies switched earlier than smaller colonies. To characterize the relation between colony size and time point of switching systematically, we measured the delay between colony formation and the time point of switching. For this series of experiments, we grew colonies in static cultures without continuous flow, because switching to collective polarization occurs faster than in the flow chamber. We found that the delay decreased with increasing initial size of the colony (Fig. 2e). Furthermore, with increasing initial cell density, the delay decreased (Fig. S5), suggesting that collective hyperpolarization is caused by depletion of nutrients or oxygen.

In conclusion, we discovered a switch in membrane polarization dynamics from independent to collective changes in membrane potential that depends on colony size and marks the transition from volume to peripheral growth.

### In the wake of the hyperpolarized shell, intracellular K_+_ levels increase and cells depolarize

After collective hyperpolarization has passed, cells depolarize slightly but significantly (Fig. 2c, 3a) and transient changes in polarization as shown in Fig. S2 disappear. In previous reports, potassium flux has been shown to be tightly correlated with changes in membrane potential (7, 9). We addressed the question whether K^+^ ions are involved in the membrane potential dynamics of gonococcal colonies. To this end, we applied the K^+^-selective ionophore valinomycin to static cultures once the hyperpolarized shell had formed. Addition of valinomycin removed hyperpolarization and the TMRM fluorescence was homogeneously reduced throughout the colony (Fig. S8). These results indicate that K^+^ ions are heavily involved in forming the characteristic pattern of membrane potential. To investigate the dynamics of intracellular K^+^, we stained *wt*^*^ colonies with TMRM and a reporter for intracellular K^+^ concentration, IPG-2 AM (Fig. 3a, b, c). We found that the IPG-2 AM fluorescence intensity strongly increased in the central region after the shell of hyperpolarized cells had passed, demonstrating that the concentration of potassium ions increased. This finding suggests that depolarization is at least partially caused by an influx of K^+^ ions. We also assessed changes in extracellular K^+^ and Na^+^ concentrations using extracellular reporters for these ions, and detected little changes. Likewise, changes in intracellular Ca^2+^ levels were not detectable.

**Fig. 3.**
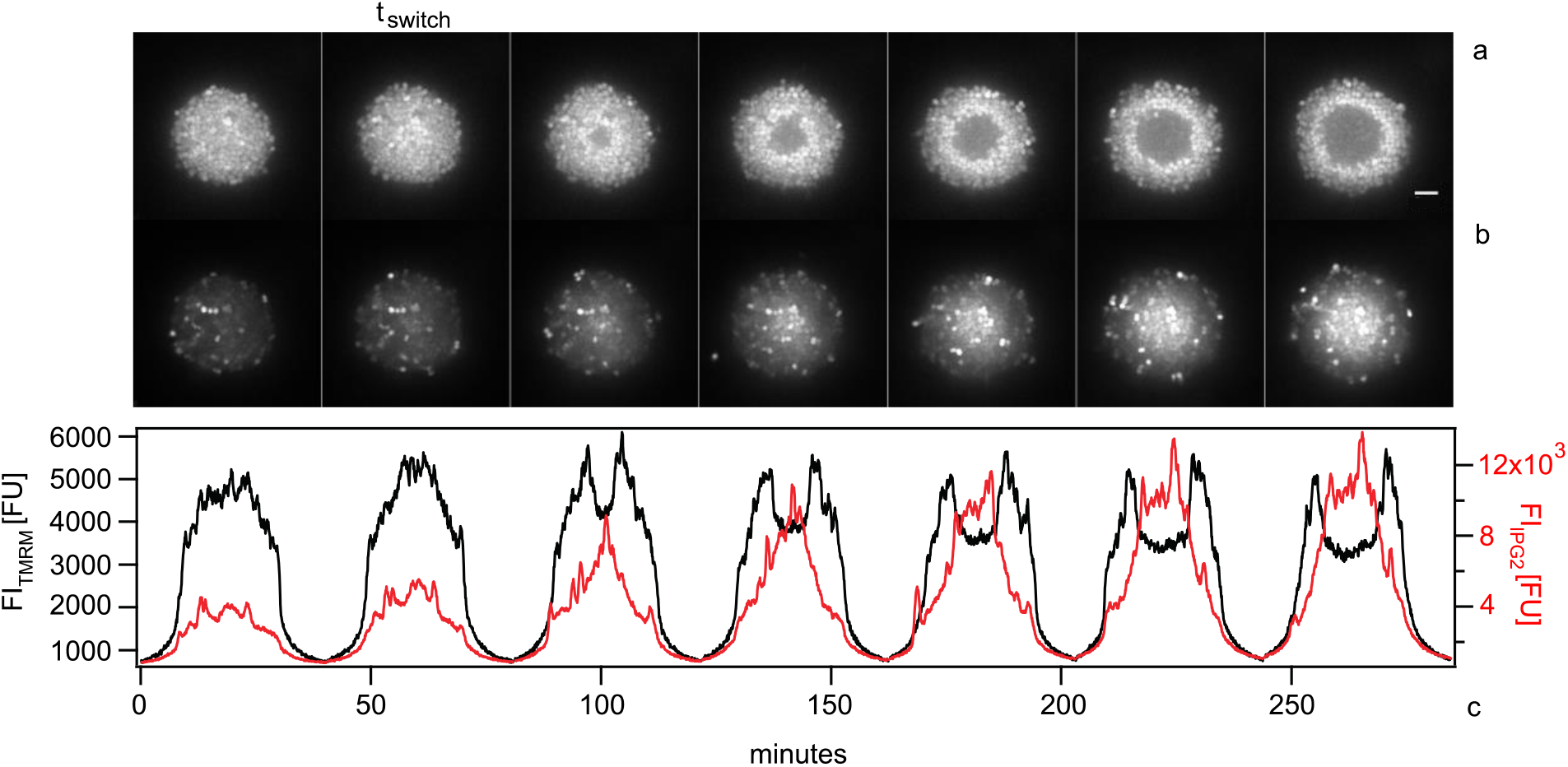
Membrane depolarization correlates with the influx of K_+_ ions. Strain *wt*^*^ (NG150), static culture. a) Time lapse of TMRM fluorescence and b) the K^+^ reporter IPG-2 AM across the equator of a colony. Scale bar: 5 μm. c) Intensity profiles along the ROI shown in a) and b).

In summary, after the hyperpolarization shell has passed, cells in its wake experience an influx of potassium ions and the membrane potential weakens.

### A reaction-diffusion model of oxygen uptake captures the dynamics of shell propagation

Next, we investigated the underlying mechanisms of shell formation and propagation. The onset of collective hyperpolarization depends on colony size (Fig. 2e), strongly suggesting that it is related to a local gradient building up within the colony. The fact that the time point of collective polarization also depends on the total concentration of cells (Fig. S5), indicates that depletion of a growth resource triggers collective hyperpolarization. The degree of oxygenation is thought to be a primary factor that influences the chemistry and physiology of a microenvironment (13, 14). Because biofilm morphology shapes the oxygen concentration profile (36), we expect that oxygen deprivation is strongest in the centre of the colony, where the hyperpolarization shell originates. Hypoxia has been shown to explain heterogeneous growth in biofilms using reaction-diffusion theory (37).

To assess whether oxygen gradient formation can explain the dynamics of the hyperpolarized shell, we first tested whether a two-dimensional reaction-diffusion model (see supplementary Methods) predicts the features of shell propagation on the observed time scales. In our model, the oxygen gradient changes with time because cells proliferate causing colony growth (Fig. 4a). We assume that the hyperpolarized shell is triggered when the oxygen concentration *c*(*t, x, y*) falls below an arbitrary critical value *c*^*^. Therefore, in our simulations, we tracked iso-concentration lines corresponding to the concentration *c*^*^, which we assume to signify the position of the hyperpolarized shell propagation. To account for growth arrest, we set the proliferation rate of cells behind these circles to zero. Furthermore, we assume that the oxygen uptake rate is reduced in this area and diminished the oxygen uptake rate of cells by a factor ε (between 0 and 1).

**Fig 4.**
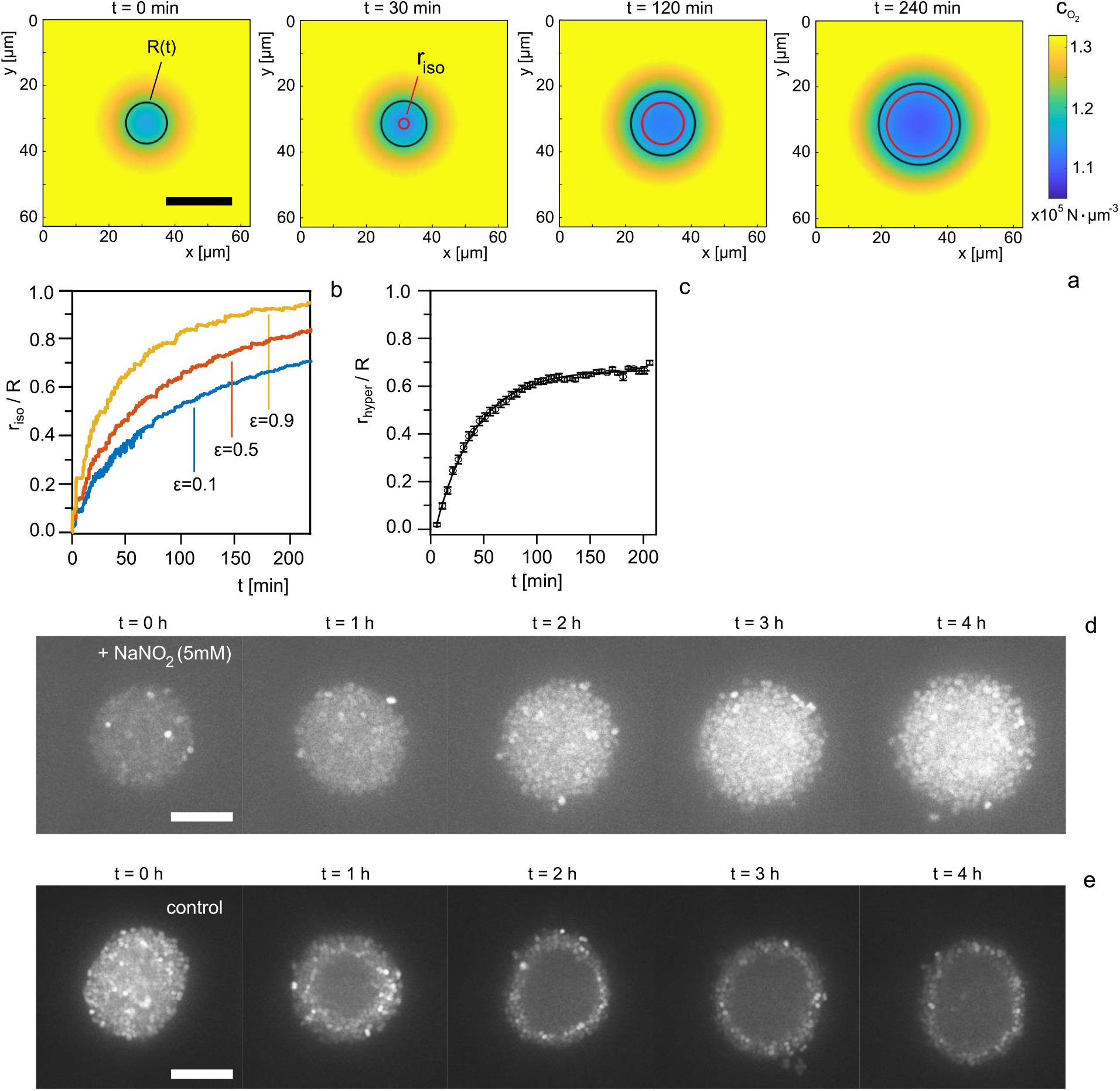
Simulation of oxygen gradients and addition of alternative electron acceptors suggest that oxygen depletion triggers collective hyperpolarization. a) Time lapse of simulated oxygen concentration field. Color code: Oxygen concentration 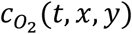. Black circles: Colony radius *R*(*t*). Red circles: Radial distance *r*_*iso*_ of the iso-concentration line at concentration *c*^*^. Scale bar: 20 μm. b) Simulated propagation of the *r*_*iso*_ circle corresponding to *c*^*^ = 0.8*c*_0_ (curves from simulations are shown for the uptake parameters ε = 0.1, 0.5, and 0.9). c) Experimental propagation of transient hyperpolarisation shells. (Means and standard errors over 20 colonies). d) Time lapse of TMRM signal for a colony (strain *wt*^*^, Ng150) in static culture supplemented with 5 mM of NaNO_2_. e) Time lapse of TMRM signal for a colony (strain *wt*^*^) in static culture without supplement. Scale bar: 10 μm.

We found that the propagation of iso-concentration circles qualitatively predicts the experimental dynamics of the hyperpolarisation shell. Circles of concentration *c*^*^ start at the centre of the colony and propagate outward at close to constant speed (Fig. 4a, b). As the circle approaches the edge of the colony, propagation captured through the relative position *r*_*iso*_/*R* slows down and stops several micrometres away from the boundary. The specific distance to the interface is governed by the free parameter *ε*; if cells at the colony centre consume less oxygen, the circle propagates more slowly and equilibrium between oxygen diffusion and uptake is reached further inside the colony (Fig. 4b). To compare the dynamics predicted by the model with the experiment, we characterized the propagation dynamics of the hyperpolarized shell by determining the normalized radius of the shell, *r*_*hyper*_/*R*, as a function of time (Fig. 4c). In agreement with the simulations, the shell propagated with a constant radial velocity early after the onset of hyperpolarization and subsequently slowed down as it reached the colony periphery. The early propagation velocity of 0.11 ± 0.01 *μm*/*min* and the characteristic propagation time *τ*_*hyper*_ = 38 ± 1 *min* obtained by an exponential fit to the data in Fig. 4c, *r*_*hyper*_/*R* = *A*(1 − *e*^−*t*/*τ*^), agree with the range of values obtained for the simulations (Fig. 4b). We note, however, that not all parameters required for the model have been characterized experimentally for *N. gonorrhoeae* (see Methods), and, therefore, we do not expect exact quantitative agreement between simulation and experiment.

If a concentration field of a growth resource is indeed involved in the onset and dynamics of the hyperpolarization shell, we expect the transiently hyperpolarized region to be broader and less defined for colonies with reduced cell number density and/or local ordering and non-spherical geometries. In this case, average radial gradients would be smeared out. Spherical colony shape and local order require the T4P retraction motor PilT (23, 25, 26, 38) and, consequently, Δ*pilT* strains form non-spherical colonies. We repeated the flow chamber experiments with Δ*pilT* colonies and found that polarization dynamics of early colonies was comparable to *wt*^*^ colonies. With time, the centres of colonies depolarized following the geometry of the aggregate, and, as predicted, there was only a weak and broad signature of collective hyperpolarization (Fig. S7 a, b). To find out whether this difference between wt* and mutant colonies was caused by lack of T4P retraction or change in colony architecture, experiments with the strain *pilT*_*WB2*_ that has strongly reduced T4P retraction activity (25), but still forms spherical colonies with local liquid-like order (25) were carried out. *pilT*_*WB2*_ colonies qualitatively behaved like wt* colonies (Fig. S7 c, d), indicating that colony morphology rather that T4P retraction was important for spatially correlated hyperpolarization.

Our model implies that the hyperpolarization shell propagates by transient hyperpolarization and not by flow of permanently hyperpolarized cells. In the following, we argue that the observed dynamics is qualitatively and quantitatively different from the radial mass flow caused by cellular proliferation. The growth-related velocity is minimal at the centre of the colony and reaches its maximum at the edge of the colony (32) in contrast to the result shown in Fig. 4c. The growth-related flow velocity is 4 to 5-fold lower compared to the propagation velocity of the hyperpolarized shell. At the same time, transient collective hyperpolarization is faster than the reshuffling of caged cells induced by pili retractions (27). Therefore, we exclude convective cell flow of permanently hyperpolarized cells (either growth or pili-driven) as the cause of shell propagation.

The propagation dynamics of the hyperpolarized shell agrees well with the dynamics of iso-concentration rings of oxygen. At this point, however, we cannot exclude that other growth resources like glucose underlie the observed polarization dynamics. Oxygen acts as the final electron acceptor in the respiratory chain of the bacteria (39). If oxygen triggers collective hyperpolarization, then addition of an alternative electron acceptor should retard or inhibit formation of the hyperpolarized shell. We investigated this point by supplementing the medium with sodium nitrite, which acts as an electron acceptor in the truncated denitrification pathway of *N. gonorrhoeae* (39-41). We found that colonies supplemented with sodium nitrite exhibit no transition to collective membrane potential dynamics over the time course of 4 hours (Fig. 4d). Conversely, control experiments consistently show collective polarization dynamics after 30 to 60 minutes (Fig. 4e). Furthermore, we exposed colonies to the oxygen scavenger 3,4-protocatechuic acid (PCA) together with its enzyme protocatechuate-dioxygenase (PCD) (33, 42) and observe a polarization pattern reminiscent of the late stages of shell propagation (Fig. S6).

In summary, we present strong evidence that the emergence of transient shell hyperpolarization and collective membrane potential dynamics is triggered by oxygen depletion along concentration gradients which form and travel outward towards the edge of the colony in radial direction.

### Tolerance to kanamycin but not to ceftriaxone is increased within the hyperpolarized shell

We investigated whether switching into the collective state affects the survivability of cells in different conditions. In particular, we addressed the emergence of antibiotic tolerance. At the single cell level, it was shown that transient hyperpolarization correlates with increased death rate (6, 8), suggesting that survivability decreases within the region enclosed by the shell of hyperpolarized cells. The state of polarization influences the bactericidal effect of aminoglycosides (8, 43), and depolarization within the shell would protect the cells. Furthermore, reduced cell growth affects the killing efficiency of certain antibiotics including β-lactams (44), predicting that cells within the hyperpolarized shell are more tolerant to this class of antibiotics than the cells outside.

First, we studied the time evolution of the fraction of dead cells within untreated colonies using the membrane-impermeable DNA stain SytoX. Once the shell of hyperpolarized cells had passed, the fraction of dead cells F increased (Fig. 5a). In particular, F inside the hyperpolarized shell was 3.5-fold higher than outside of the shell (Fig. 5b). This finding is consistent with earlier studies showing that the death rate of individual cells that underwent transient hyperpolarization was higher than the death rate of cells without hyperpolarization (6, 8).

**Fig. 5.**
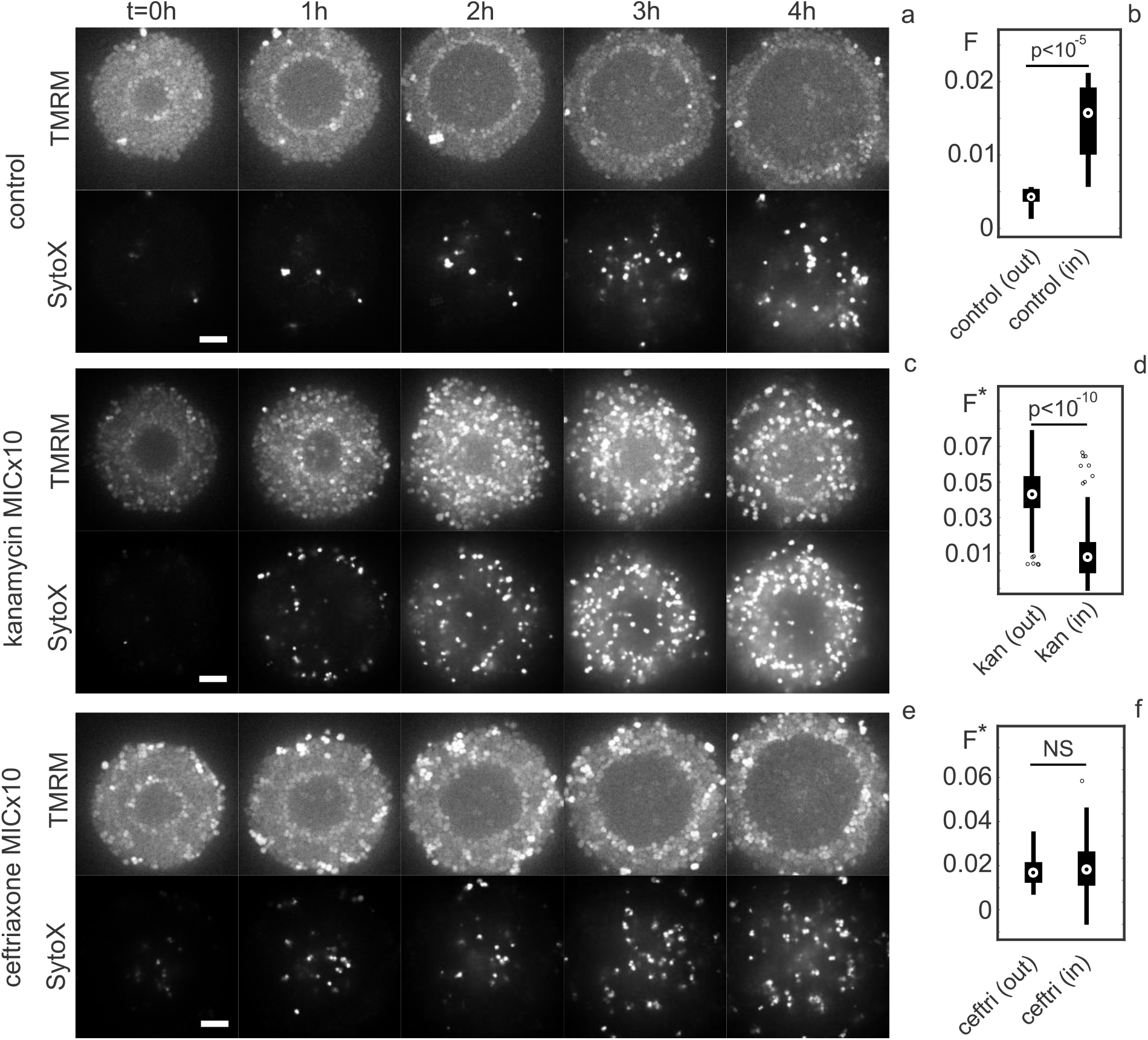
Dynamics of membrane potential and fraction of dead cells *F* in colonies after 30 min of antibiotic treatment. Strain *wt*^*^ (NG150), flow chamber. a), c), e) Time lapse over 4 hours of TMRM and SytoX fluorescence after respectively: no treatment, kanamycin treatment (MIC×10, 200 μ*g*/*ml*), and ceftriaxone treatment (MIC×10, 0.04 μ*g*/*ml*). Scale bar: 5 μm. b), d), f) Fraction of dead cells *F* in the colonies 3-4 hours later for both the region unaffected by the hyperpolarisation shell (out) and the region affected (in) after respectively no treatment, kanamycin treatment, and ceftriaxone treatment. In the treated cases, *F*^*^ = *F*−< *F*_*control*_ >. 60 colonies evaluated for each condition. Bottom and top edges of the box indicate the 25th and 75th percentiles, respectively. Whiskers extend to the most extreme data points not considered outliers (marked as individual points). Central mark indicates the median of the distribution.

Next, we addressed the questions how treatment with the aminoglycoside kanamycin affects membrane potential dynamics and how the pre-formed polarization pattern impacts kanamycin tolerance. Two-hour old colonies were treated with kanamycin at 10-fold MIC for 30 minutes. After removal of the antibiotics, we characterized the dynamics of the membrane potential and the fraction of dead cells. Transient treatment with kanamycin caused a gradual increase in polarization throughout the colony, typically starting at the edge of the colony and moving inward (Fig. 5c). Surprisingly, for colonies which had already formed a shell of hyperpolarized cells, kanamycin inhibited its radial progression and often the shell slowly travelled back towards the centre of the colony, where it disappeared if it reached the centre. Similar dynamics were observed under azithromycin treatment (Fig. S9). To find out whether tolerance increased in the wake of the hyperpolarized shell, we compared the regions of the colonies that resided inside the hyperpolarized shell during antibiotic treatment with the outside region. The fractions of dead cells were determined 3 to 4 hours after treatment, because cell death is detectable with SytoX at a delay of several hours (29). To account for the fractions of dead cells in the absence of antibiotics, we subtracted the respective values (Fig. 5b). We found that the fraction of dead cells within the shell was 4-fold lower than outside (Fig. 5c, d), indicating that depolarization increases tolerance against kanamycin despite the preceding transient hyperpolarization.

Finally, we treated the colonies with the *β*-lactam antibiotic ceftriaxone using the same protocol described for kanamycin. In contrast to the protein synthesis inhibitors described above, ceftriaxone did not affect the membrane potential (Fig. 5e). After subtraction of the fraction of dead cells without treatment, the fraction of dead cells was homogeneous throughout the colony (Fig. 5e, f), showing that the pattern of membrane potential and the associated growth rate reduction at the centre did not affect tolerance against ceftriaxone.

In summary, kanamycin treatment affects the polarization pattern of the colony whereas ceftriaxone has no effect. Within the hyperpolarized sphere, cells were more tolerant against kanamycin but not against ceftriaxone.

## Discussion

Transient hyperpolarization has been reported at the single cell level for *E. coli* and *B. subtilis* (4, 6, 8). Importantly, hyperpolarization was uncorrelated between neighbouring cells in these studies. Similarly, the membrane potential dynamics of freshly assembled gonococcal colonies studied here shows no nearest neighbour correlation. This suggests that gonococci, while being part of multicellular colonies, still behave like planktonic cells in the first few hours after assembly. By contrast to previous reports, we reveal a switch from independent to correlated hyperpolarization occurring once the colony is large enough.

Our results are most consistent with the following scenario. Governed by T4P motility and T4P-mediated cell-to-cell adhesion, gonococci assemble into colonies by T4P activity within minutes (23). During the early stage of this colony, gonococcal behaviour remains uncorrelated and colony formation is fully reversible (19). Within the colony, a gradient of oxygen builds up. When a critical concentration of oxygen is reached, cells hyperpolarize. Once the oxygen concentration falls below this critical concentration, cells depolarize, are growth-inhibited, and consume less oxygen. Since the colony continues to grow at the surface, the critical iso-concentration shell (signified experimentally as a hyperpolarized shell) propagates towards the periphery of the colony. Depending on the oxygen consumption rates, the shell of hyperpolarized cells becomes stationary at a specific distance away from the edge of the colony. An analytical solution of our reaction-diffusion model (see Supplementary Information) predicts that this distance is related to the critical O_2_ concentration *c*^*^, the concentration at the surface of the colony, *c*_0_, and the concentration *c*_*c*_ at the centre of the colony, 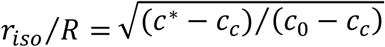. In the flow chamber, we find *r*_*hyper*_/*R* ≈ 0.7, which entails that the critical concentration is in the range 0.5*c*_0_ < *c*^*^ < *c*_0_. Under static growth conditions, we observed that the hyperpolarized shell is closer to the edge of the colony (see e.g. Fig. 4e), in agreement with the O_2_ concentration at the surface being lower than under continuous flow. It is also interesting to note that treatment with the protein synthesis inhibitors kanamycin and azithromycin halts or reverses the propagation of the hyperpolarized shell. As the antibiotics inhibit the growth of the colony and potentially reduce the consumption rate of oxygen, our model would predict this observation.

While, we can clearly demonstrate that collective and transient hyperpolarization signifies the transition of volume to surface growth and changes in the state of antibiotic tolerance, the molecular mechanism of transient hyperpolarization remains unclear. According to our model, the O_2_ concentration range that causes hyperpolarization is narrow. Hyperpolarization correlated with growth arrest or increased mortality has been reported in various studies (6-8). One study identifies a cause for hyperpolarization; under aminoglycoside treatment the inhibition of ribosomes frees ATP which causes the ATP synthase to work in ATP hydrolysis direction, increasing the proton gradient (8). In our system, we rather expect the ATP levels to be reduced as a consequence of reduced proton gradient under low oxygen conditions. It will be very interesting to investigate the mechanism of transient hyperpolarization in future work.

We expected that reduced polarization and growth rate made gonococci more tolerant against aminoglycosides and β-lactams, respectively. The absolute decrease in polarization between the outside and the inside of the hyperpolarized shell amounts to a few millivolts only. Yet, we clearly observed that the fraction of dead cells is lower within than outside of the shell after transient kanamycin treatment, indicating that cells are immediately protected against this aminoglycoside. By contrast, growth arrest within the hyperpolarized shell had no influence on tolerance against the β-lactam ceftriaxone. A previous study showed that tolerance quantitatively correlates with growth rate (44). In that study, growth rates were varied by different but constant growth environments. The protective effect of growth inhibition may not occur instantaneously after reduction of growth rate explaining why gonococci were not protected against ceftriaxone despite growth inhibition.

In summary, we reveal a switch from uncorrelated single cell fluctuations to collective dynamics of membrane polarization in gonococcal colonies and provide strong evidence that the local oxygen gradient governs this transition. Further, we find that a traveling shell of hyperpolarized cells demarks the differentiation into two subpopulations with different growth rates and different levels of antibiotic tolerance. In future studies it will be important to find the underlying mechanism of transient hyperpolarization. We anticipate that studying dynamical polarization patterns in developing biofilms will provide novel insights into the mechanisms of biofilm differentiation in various bacterial species.

## Supporting information

Movie S1. Typical example of colony with blinking cell. Movie corresponds to Fig. S2a. dt = 1 min.

Movie S2. Three-dimensional reconstruction of formation and propagation of the hyperpolarized shell. dt = 5 min.

Movie S3. Typical example of onset of collective hyperpolarization for colonies with different initial sizes. dt = 5 min.

## Acknowledgements

We thank the Maier lab for stimulating discussions. This work was supported by the Deutsche Forschungsgemeinschaft through grant MA3898 and the Center for Molecular Medicine Cologne.

