## Supplementary Methods, Figures, Tables for "Collective polarization dynamics in gonococcal colonies"

### Materials and methods

**Growth media and bacterial strains.** Media composition, assay preparation and setup follow (27). Gonococcal base agar was made from 10 g/L dehydrated agar (BD Biosciences, Bedford, MA), 5 g/L NaCl (Roth, Darmstadt, Germany), 4 g/L K2HPO4 (Roth), 1 g/L KH2PO4 (Roth), 15 g/L Proteose Peptone No. 3 (BD Biosciences), 0.5 g/L soluble starch (Sigma-Aldrich, St. Louis, MO), and supplemented with 1% IsoVitaleX (IVX): 1 g/L D-glucose (Roth), 0.1 g/L L-glutamine (Roth), 0.289 g/L L-cysteine-HCL × H2O (Roth), 1 mg/L thiamine pyrophosphate (Sigma-Aldrich), 0.2mg/LFe(NO3 )3 (Sigma-Aldrich), 0.03 mg/L thiamine HCl (Roth), 0.13 mg/L 4-aminobenzoic acid (Sigma-Aldrich), 2.5 mg/L β-nicotinamide adenine dinucleotide (Roth), and 0.1 mg/L vitamin B12 (Sigma-Aldrich). GC medium is identical to the base agar composition but lacks agar and starch. Employed bacterial strains (Table 1) are derivatives of strain *N. gonorrhoeae* MS11.

**Culture preparation for experiments in the flow chamber and under static conditions.** For the microscopy experiments we used two different setups. First, bacteria were grown in flow chambers. This condition ensures continuous flow of medium and approximately constant supply of growth resources. Second, bacteria were grown in static culture. This condition was used when fast depletion of growth resources was desired or supplements were added during the experiment. All experiments follow the same preparation protocol: Overnight cultures of the respective strain were suspended in fresh GC + IVX medium at $OD_{600}$ 0.1 unless noted otherwise. Under addition of 1% v/v ddH2O, the obtained bacterial solution was then incubated in a shaker at $37^o\ C$ and 5% $CO_2$ for 30 minutes to 1 hour, which allowed colonies to first disassemble and then reform freshly.

Flow chamber experiments: Cells were loaded into a microfluidic flow chamber (Ibidi Luer 0.8 mm channel height + Ibitreat) which was previously coated with Poly-L-Lysine (Sigma, Cat. No. P4832, 50 μg/mL) overnight. The flow chamber was continuously supplemented with medium at a temperature of $37^o\ C$ using a peristaltic pump (model 205U;Watson Marlow, Falmouth, United Kingdom) operating at 2 rpm. To visualize the membrane potential, TMRM (Sigma-Aldrich) was added to the supplied medium at a concentration of 0.1 $\mu$M (33).

Static condition experiments: 300 $\mu L$ of the bacterial suspension at an OD of 0.1 were pipetted into a $\mu$-slide 8 Well (Ibidi + Ibitreat), which was coated with Poly-L-Lysine (Sigma, Cat. No. P4832, 50 μg/mL) over the previous night. Together with the bacterial suspension, and depending on the experiment, we added 3 $\mu$M valinomycin (Sigma-Aldrich); 5 mM sodium nitrite NaNO2 (Roth); or 20 $\mu g$/mL IPG2-AM (Ion BioSciences). Oxygen scavenger system (protocatechuic acid, PCA, and protocatechuate 3,4-dioxygenase, PCD, both from Sigma-Aldrich) preparation and concentration was taken from (42).

**Confocal imaging.** For imaging, colonies of various sizes and diameters were randomly chosen and imaged using an inverted microscope (Ti-E Nikon) equipped with a CSU-X1 (Yokogawa) spinning disk unit, a 100x CFI Apo Tirf objective (Nikon), an EMCCD camera (iXon 897 X-11662, Andor) and 488 nm and 561 nm excitation lasers. If not stated otherwise, colonies were imaged at their equator once every minute (acquisition time 100 ms) in the brightfield, green, and red channel (laser power 3 % and 2 %, respectively).

**3D movie of hyperpolarized shell progression.** 3D movies were obtained by recording multiple colonies inside the flow chamber every 5 minutes (strain NG194, $wt\ green$, acquisition time: 50 ms) for a total time interval of 2 hours. For every imaging step, we recorded a z-stack of 10 µm with a z-spacing of 0.2 µm between each image close to the bottom of the microscope coverslip. 488 nm and 561 nm excitation lasers were set to 5 % and 10 % intensity, respectively. In addition, we acquired a brightfield image for every time point. Finally, 3D volume renderings and movies of hyperpolarization shell formation within colonies were created with NIS-Elements Ar Ver. 5.02 (Build 1271, Nikon).

**Characterization of antibiotic tolerance and fraction of dead cells.** Colonies ($wt^*$, NG150) were incubated for ~120 minutes to allow the formation of the hyperpolarized shell within a subset of colonies. Subsequently, colonies were treated with medium containing kanamycin, ceftriaxone, or azithromycin at a concentration corresponding to 10x the MIC for kanamycin and ceftriaxone (200 $\mu g$/mL and $0.04\,\mu g$/mL, respectively), and 100x the MIC for azithromycin ($0.64\,\mu g$/mL) for 30 minutes. Under control conditions, colonies were treated with medium which was supplemented with an equal volume of dd$H_2O$ instead. After the treatment was stopped, colonies were again supplemented with medium lacking any antibiotics but containing SytoX (Invitrogen) at a concentration of 0.2 $\mu$M. During this process, we acquired a z-stack of each colony every 10 minutes. The stack covered an area of 5 µm around the equatorial plane of the colonies, with a z-spacing of 0.5 µm between individual slices. Imaging was continued for 4 hours beyond the treatment phase. In addition, brightfield images were acquired. The fraction of dead cells inside of colonies was then determined by tracking the green SytoX signal (Trackmate).

**Determination of single cell potential.** The cationic dye TMRM partitions between the cytoplasm (concentration $c_{in}$) and the extracellular space (concentration $c_{out}$) depending on the electrical potential $V_m$ across the inner membrane (4)

$$V_m = \frac{RT}{zF} \ln\left(\frac{c_{out}}{c_{in}}\right) \quad (1)$$

Here, $R$ is the gas constant, $T$ is the absolute temperature, $F$ is the Faraday constant, and $z$ is the valence of the dye $(z = +1)$. Concentrations $c_{in,out}$ are proportional to the TMRM fluorescence intensities inside of cells and outside of the colonies, respectively. To derive $V_m$ from the fluorescence intensities, we proceeded as follows.

From the recordings, single cell membrane potentials were obtained by first finding cell positions for each frame through the sfGFP signal using the Fiji plugin Trackmate. We then averaged the TMRM signal (red channel) over the area of the cells at these positions. Since TMRM molecules partition between the inside and outside of the cell depending on the presence of negative charges, we additionally measured the TMRM signal far away from the colonies. For both, we subtracted the contribution of the camera noise, which we quantified from images in the absence of cells and TMRM. Membrane potential values were then calculated according to equation (1), with $T = 310K$.

We note that in this study we have not corrected for effects of diffraction. In a previous study at the single cell level, we did account for this effect and measured slightly (~ 10 %) more negative membrane potential (33). The difference may be accounted to the fact that we did not correct for diffraction here, or to the fact that cells depolarize directly after colony formation. Importantly, the difference is small and does not affect the major conclusions of this study.

**Localized growth rate before and behind the hyperpolarization shell.** To show that growth rates decline after passage of the hyperpolarized shell, but not prior to it, we devised a simplified binary colony growth model where we assume that the cell growth rate $\lambda = \lambda_0$ is constant prior to the passage of the shell, and $\lambda = 0$ afterwards. We start from the differential equation for the number of cells $N_r$ inside spherical shells at a distance $r$ from the centre of mass of the spherical colony: $\dot{N}_r = \lambda(r,t)N_r$. With $N_r = \phi V_r$, where $\phi$ is the cell number density of the shell and $V_r$ its volume, we can write the temporal change in total colony volume as $\dot{V} = \int_0^R \lambda r^2 dr$ ($R$: colony radius). Using $\dot{V} = 4\pi \dot{R} R^2$, and splitting the integral in a part behind ($r < r_{hyper}$) and before ($r > r_{hyper}$) the hyperpolarized shell, we obtain a relation between colony radius $R$ and the instantaneous radial shell position $r_{hyper}$

$$R^2\left(R - \frac{3\dot{R}}{\lambda_0}\right) = r_{hyper}^3$$

Before the hyperpolarized shell forms ($t < t_{switch}$), we expect the left-hand side to be equal to zero. Afterwards, the LHS evolves as the third power of $r_{hyper}$. This relation is in good agreement with experimentally obtained values of $R, r_{hyper}$, see Fig. S6 (15 colonies evaluated). Additionally, heterogeneous growth rates in the colonies several hours after incubation are in agreement with previous findings (32).

### Theoretical modelling

**Reaction-diffusion model for shell propagation dynamics.** We numerically solved a two-dimensional reaction-diffusion model for the concentration field of oxygen (45)

$$\partial_t c = -\kappa(c)\varrho + D\Delta c$$

Here, $c \coloneqq c(t,x,y)$ is the local concentration of $O_2$ inside and outside of the colony ($r = \sqrt{x^2+y^2}$: distance to colony centre-of-mass), $\kappa(c)$ is the oxygen consumption rate by each cell, $\varrho$ is the local cell number density, and $D$ is the diffusion coefficient of oxygen in the solvent. To account for both uptake- and diffusion-limited regimes, we used the Michaelis-Menten form $\kappa(c) = \kappa_0 \frac{c}{c+K_M}$, where $K_M$ is the Michaelis-Menten constant of the oxygen uptake reaction, and $\kappa_0$ is the upper limit of oxygen uptake per cell. As fixed parameters, we used the Michaelis constant determined for *Bacillus licheniformis*, $K_M = 2.5 \times 10^3\, molecules \cdot \mu m^{-3}$ (46), $c_{solvent} \equiv c_0 = 1.3 \times 10^5\, molecules \cdot \mu m^{-3}$ (33). In bulk solvent, the diffusion coefficient of oxygen is $D_{O_2} = 1.5 \times 10^5 \mu m^2 \cdot min^{-1}$ (47). To account for hindered diffusion in the colony, we set the oxygen diffusivity inside the colony to 1/10th this value. Finally, the product of the maximal oxygen consumption rate and the cellular density in colonies is $\kappa_0\varrho = 45 \times 10^5\, molecules \cdot \mu m^{-3}\, min^{-1}$ (25, 33). The cell density was assumed to be constant inside a circle of radius $R$, where $R$ is the radius of the colony, and zero outside. To mimic nutrient supply in the flow chamber, we set $c = c_0$ for $r = R(t) + 5\,\mu m$. We solved the model using a 2D DuFort-Frankel scheme in Matlab on a grid of size 128x128 in simulation units, corresponding to a window of 64x64 $\mu m^2$ ($ds = 0.5\,\mu m$), with $c(t = 0, x, y) \coloneqq c_0$ (timestep $\Delta t = 10^{-3}$ minutes). After $10^4$ steps (10 minutes), we obtained a steady-state concentration profile $c(t,x,y) \coloneqq c(r) \equiv c_{eq}(r)$. At that moment, we allowed cells to proliferate with growth rate $\lambda = 0.01\, min^{-1}$ (32) for 300 minutes and chose an adequate concentration $c^* = 0.8 \times c_0$, slightly smaller than $c_{eq}(0)$. From that moment, in regions where the concentration was lower, the growth rate was set to zero. The colony radius evolved as $\dot{R}(t) = \frac{\lambda}{3}\left(R(t) - \frac{r^3(t,c^*)}{R^2(t)}\right)$, where $r(t,c^*)$ is the distance of the iso-concentration circle to the colony centre-of-mass. Additionally, we allowed the oxygen uptake rate in the region $r < r(t,c^*)$ to diminish by a factor $\varepsilon \in [0,1]$. To test the robustness of our system and results, we varied several key parameters (including oxygen diffusivity, distance of boundary condition), and found that variations had little to no qualitative impact on the results.

An analytical steady-state solution to the reaction-diffusion equation in terms of a Maclaurin series was derived in (48). To quadratic order in the centre-of-mass distance $r$, the solution reads $c(r) = c_c + \frac{\kappa_0 \varrho c_c r^2}{3! D(c_c + K_M)}$ , where $c_c$ is the (time-dependent) oxygen concentration at the centre of the colony. From there, we obtain the relation $\frac{r_{iso}}{R} = \left[\frac{c^* - c_c}{c_0 - c_c}\right]^{0.5}$ under the assumption that $c_0 \approx c(R)$. Because $c_c$ is larger than zero and cannot exceed the boundary value $c_0$, the critical concentration $c^*$ has range $0.5c_0 < c^* < c_0$.

### Supplementary Figures

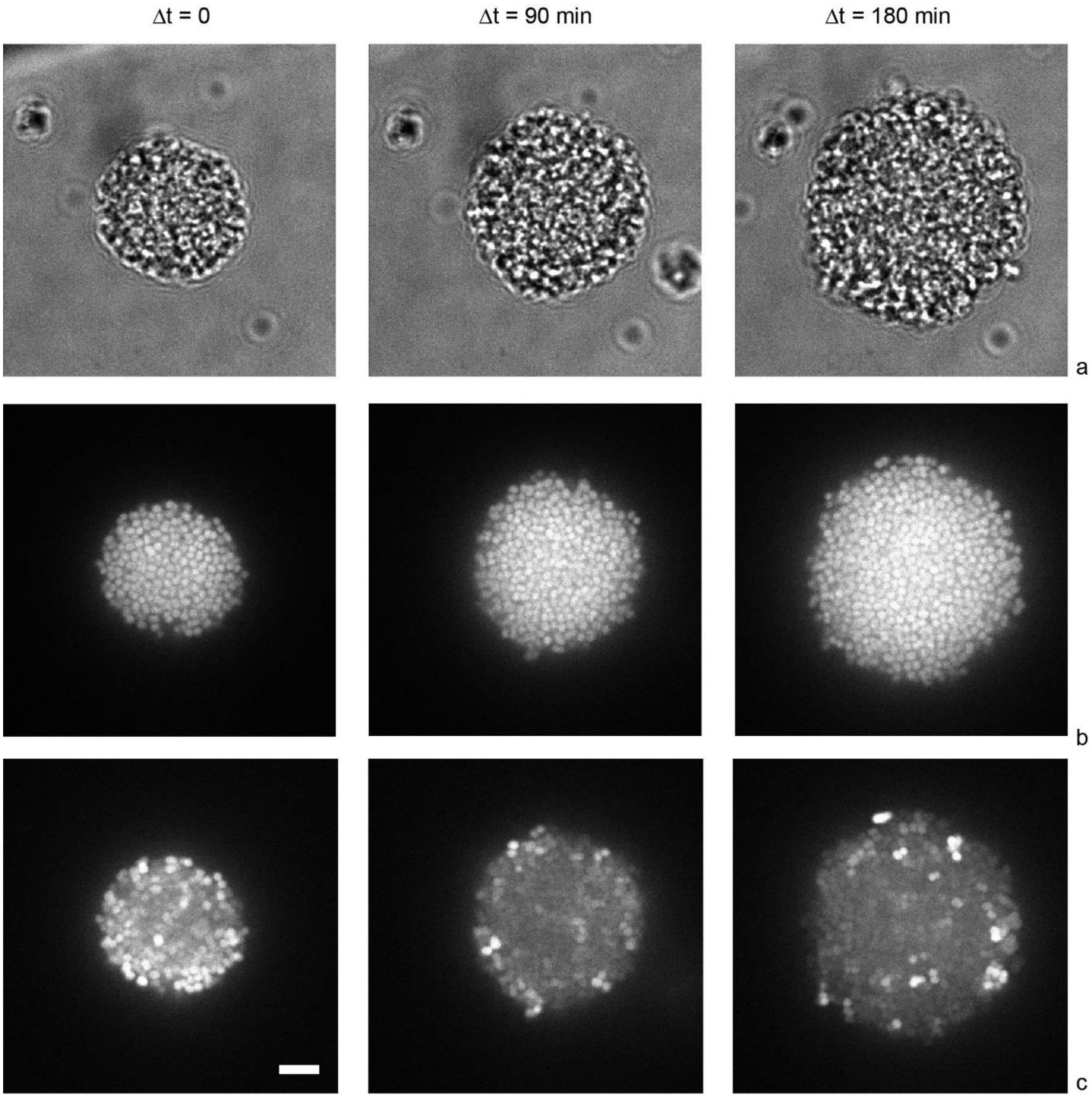


**Fig. S1 Comparison between TMRM and sfGFP fluorescence in colonies.** Strain $wt\ green$ (NG194) in flow chamber. a) Brightfield images of colonies imaged at the indicated time points. b) sfGFP fluorescence. c) TMRM fluorescence. Scale bar: 5 µm.

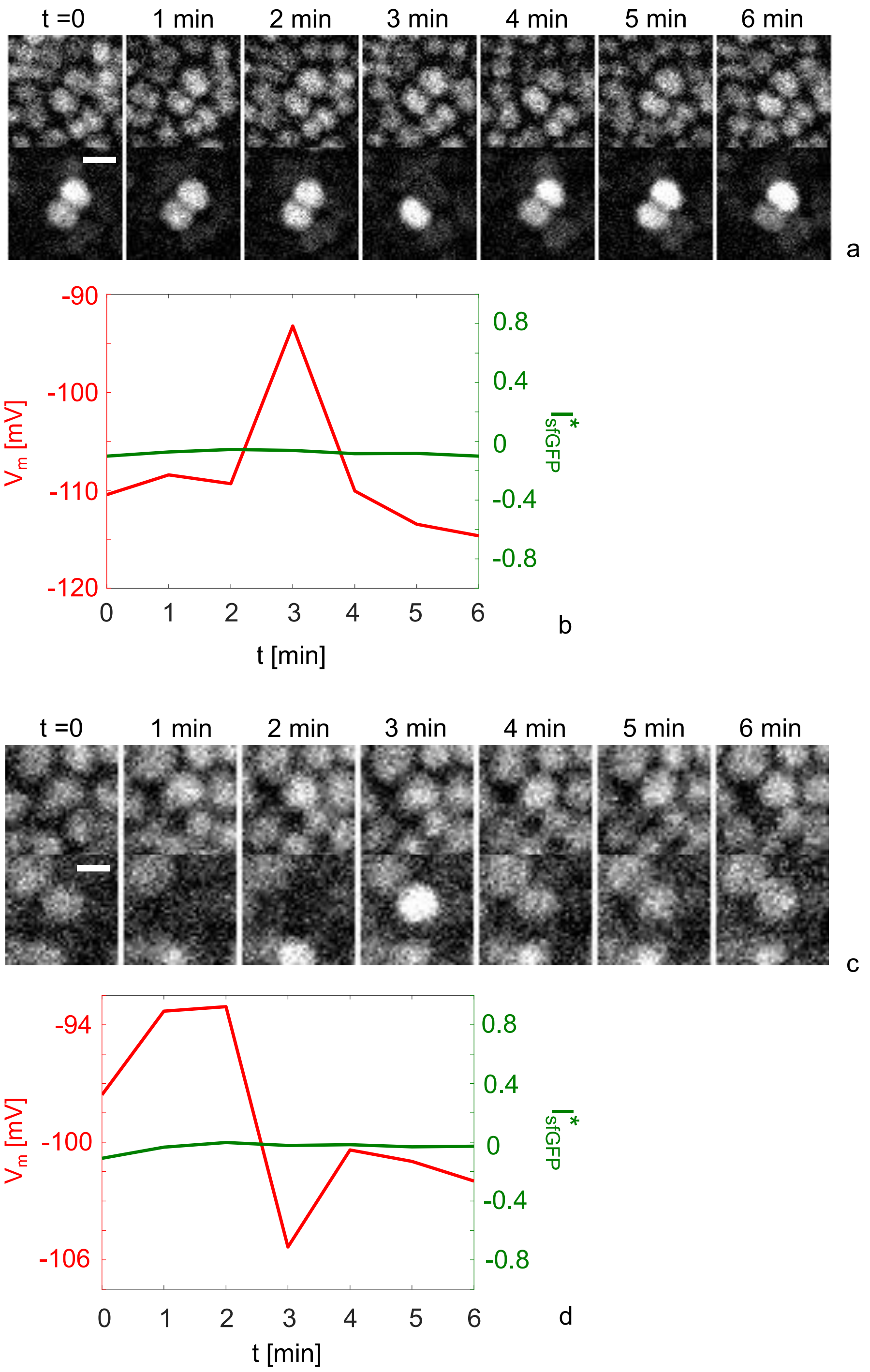


**Fig. S2: Transient depolarization and hyperpolarization events of single cells occur on the time scale of minutes and are uncorrelated between neighbours.** Strain $wt\ green$, NG194, flow chamber. a) sfGFP (top) and TMRM (bottom) signal time lapse during a transient depolarization event. $\Delta t$ = 1 min. b) Membrane potential (left axis) and normalized sfGFP-signal (right axis) $I^{*}_{sfGFP} = (I_{sfGFP} - < I_{sfGFP} >_{cells}) \,/ < I_{sfGFP} >_{cells}$ for the time lapse of a). c) sfGFP (top) and TMRM (bottom) signal time lapse during a transient hyperpolarization event. $\Delta t$ = 1 min. d) Membrane potential (left axis) and normalized sfGFP-signal (right axis) for the time lapse of c). Scale bar: 1 $\mu$m

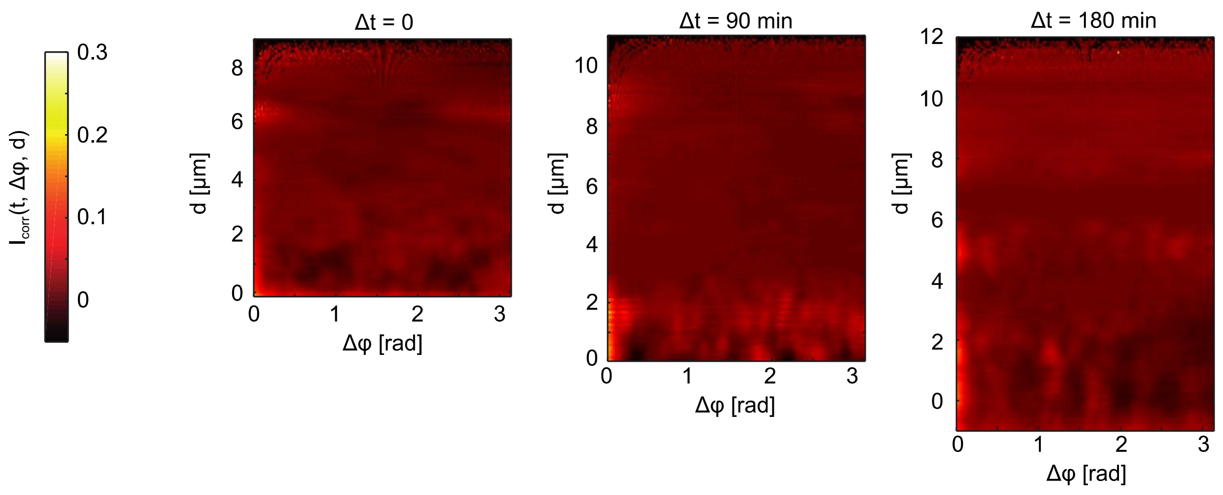


**Fig. S3 Further example of angular correlation of the intensity fluctuations within the colony prior to switching.**

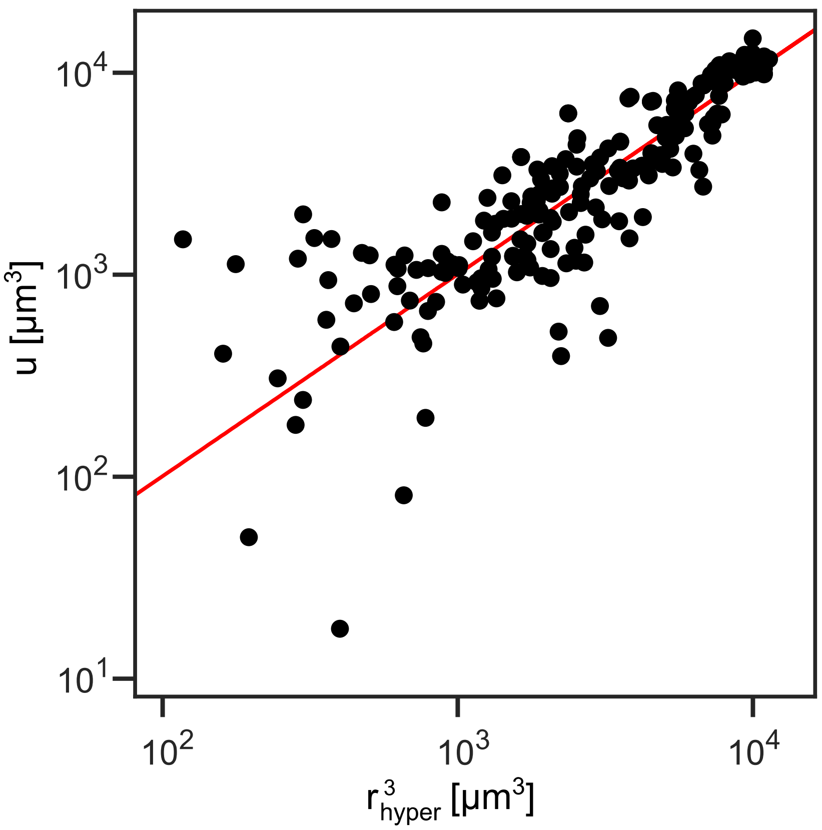


**Fig. S4 Relation between the cubed hyperpolarized shell radius $\boldsymbol{r^3_{hyper}}$ and the colony radius factor $\boldsymbol{u \equiv R^2\left(R - \frac{3\dot{R}}{\lambda_0}\right)}$.** Strain *wt green*, NG194, flow chamber. Each point represents one measurement of both variables (15 colonies evaluated). Red line: fit to power function $y = x^b$ ($b = 1.006 \pm 0.004, R^2 = 0.86$).

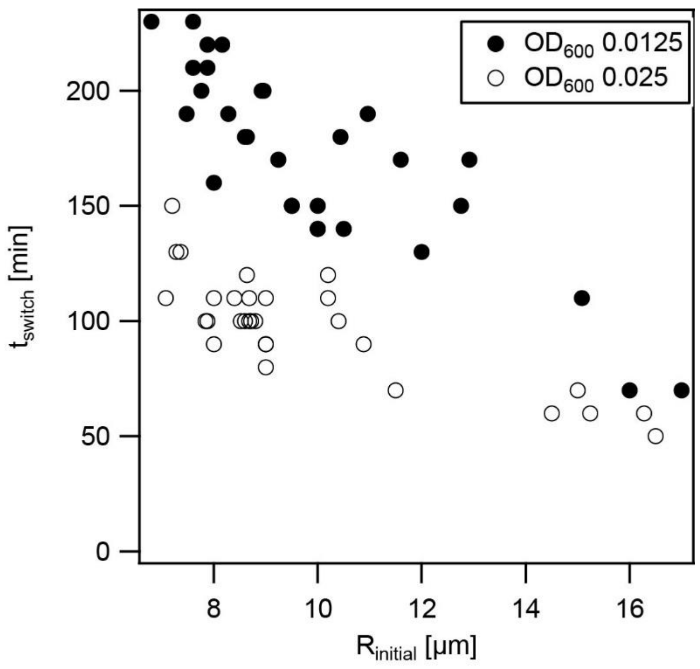


**Fig. S5 The time point of switching to collective behaviour depends on initial colony size and cellular concentration.** Strain *wt green*, NG194, static culture. The inoculation densities were $OD_{600}$ 0.0125 (filled circles) and $OD_{600}$ 0.025 (open circles), respectively.

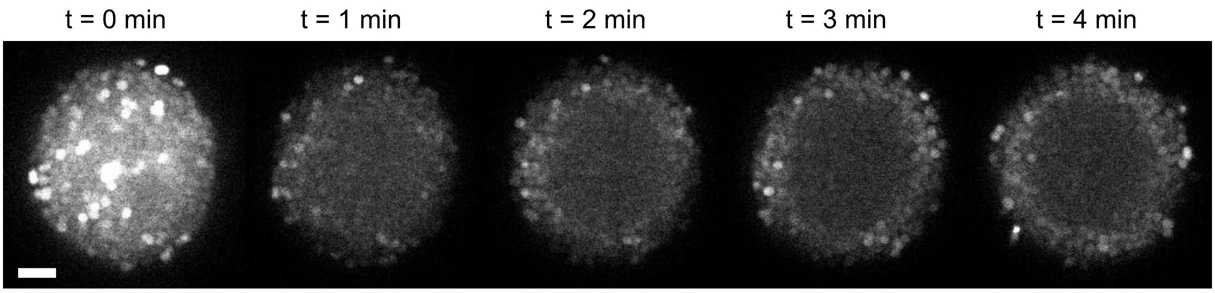


**Fig. S6 Addition of an oxygen scavenger rapidly generate the polarization pattern of large colonies.** Strain $wt^*$ (NG150) static culture. Time lapse of TMRM signal of a colony supplemented with the oxygen scavenging system PCA+PCD. Shortly after addition (t = 1 minute), the cells depolarize. After 5 minutes, the membrane potential is similar to the one found after shell passage. Scale bar: 5 $\mu$m.

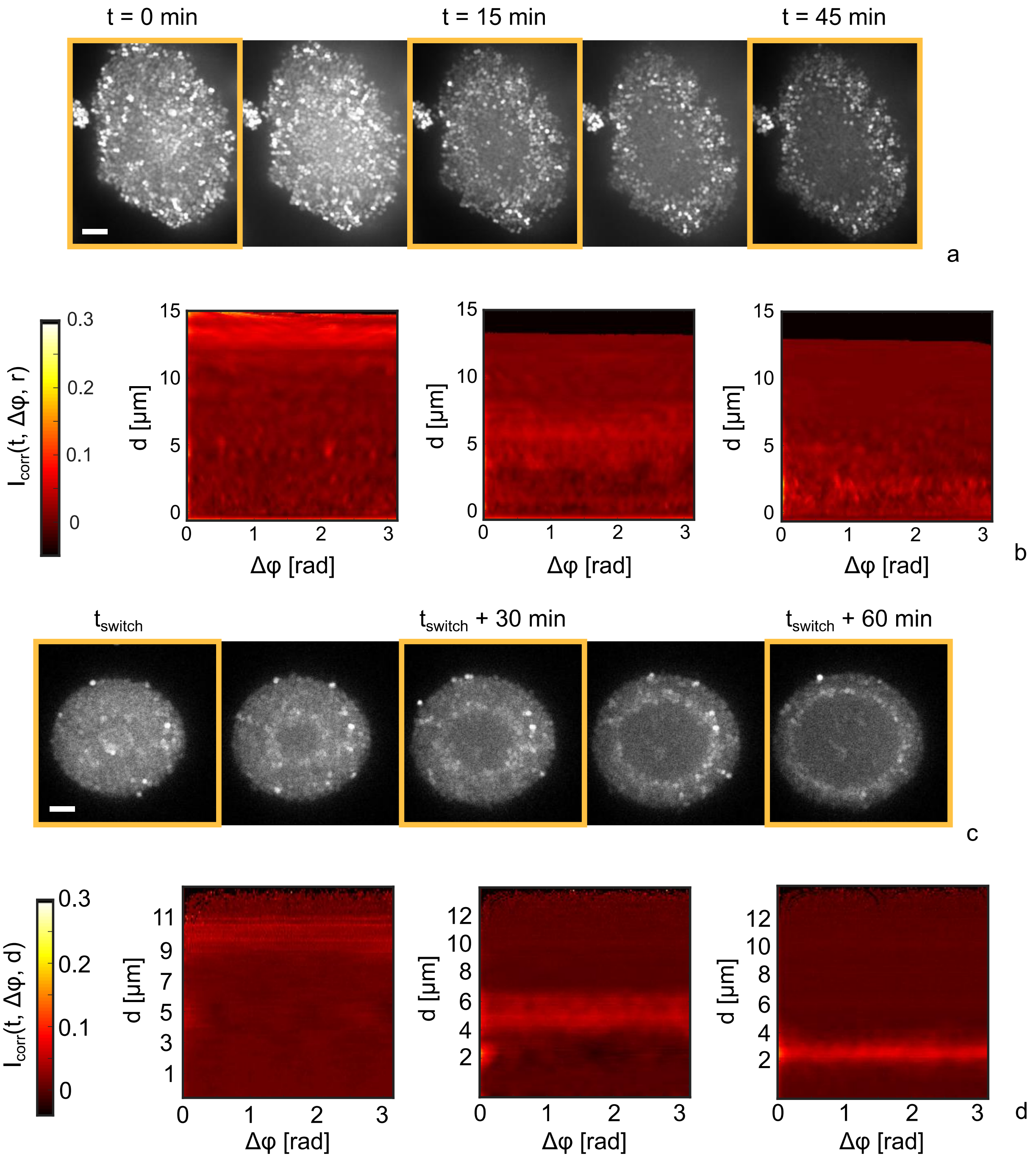


**Fig. S7 Membrane potential dynamics for a retraction-deficient and retraction-reduced strain.** Flow chamber. a) Typical time lapse of TMRM fluorescence of the T4P retraction deficient strain *ΔpilT* (NG231). Scale bar: 5 $\mu$m. b) Angular correlation of the intensity fluctuations as a function of radial position for strain *ΔpilT* at time points t=0, 15 min, 45 min (t=0 min: start of colony centre depolarization). Images correspond to timeframes in a) marked by orange boxes. c) Typical time lapse of TMRM fluorescence of strain $pilT_{WB2}$ (NG176). Scale bar: 5 $\mu$m. b) Angular correlation of the intensity fluctuations as a function of radial position for strain $pilT_{WB2}$. Images correspond to timeframes in c) marked by orange boxes.

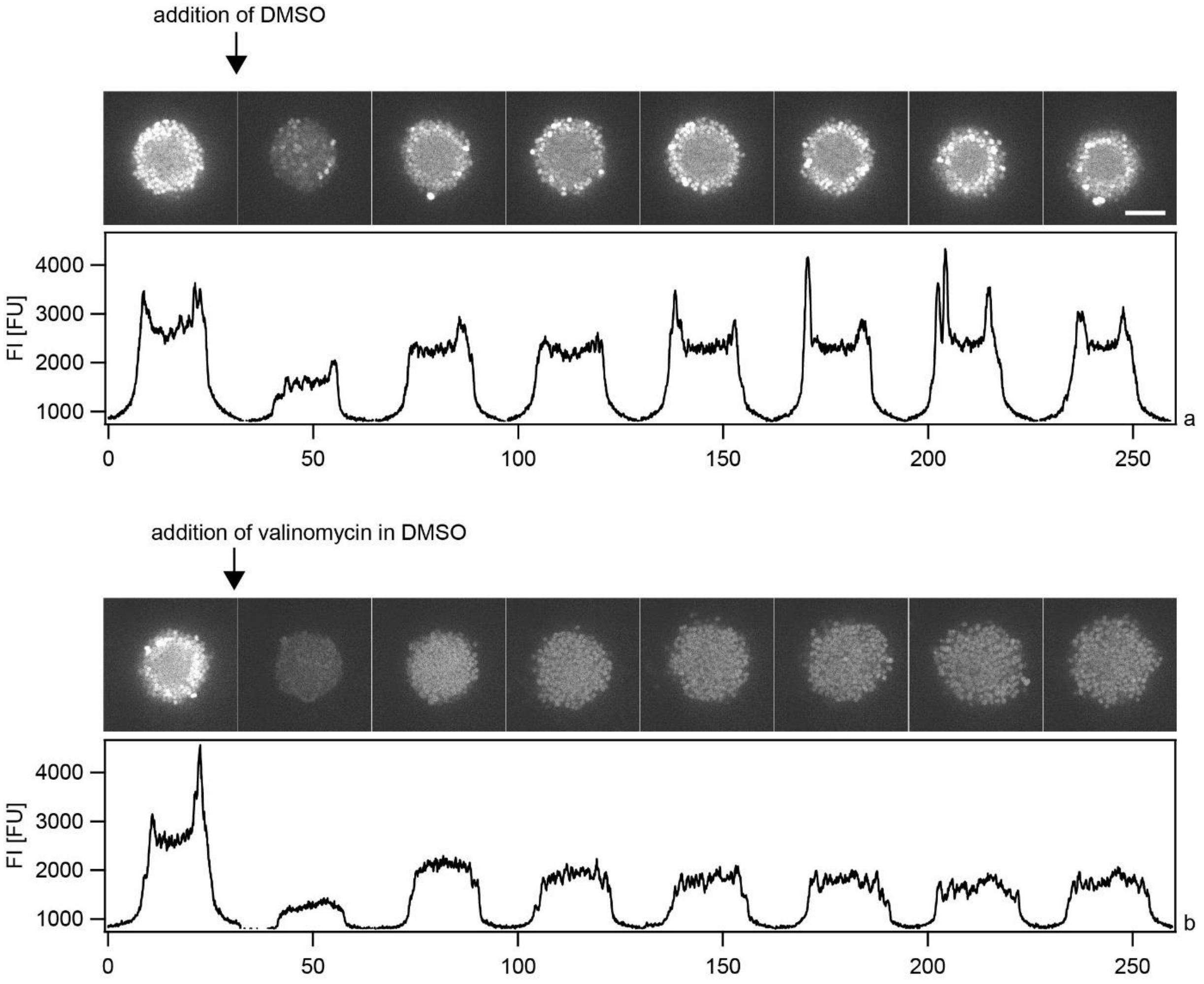


**Fig. S8 Addition of the $K^+$ selective ionophore homogeneously depletes the membrane potential**. Strain $wt^*$ (NG150), static culture. After the shell of hyperpolarized cells had travelled to the edge of the colony, valinomycin was added. a) Control. DMSO was added at the indicated time point. Immediately after addition of DMSO, cells depolarized transiently. Within 5 min, the shell had re-formed and the membrane potential pattern was restored. b) Valinomycin in DMSO was added to a final concentration of 3 µM. After the initial depletion, the membrane potential was only partially restored and no hyperpolarization was evident. Colonies started to disassemble. Δt = 5 min. Scale bar: 10 µm.

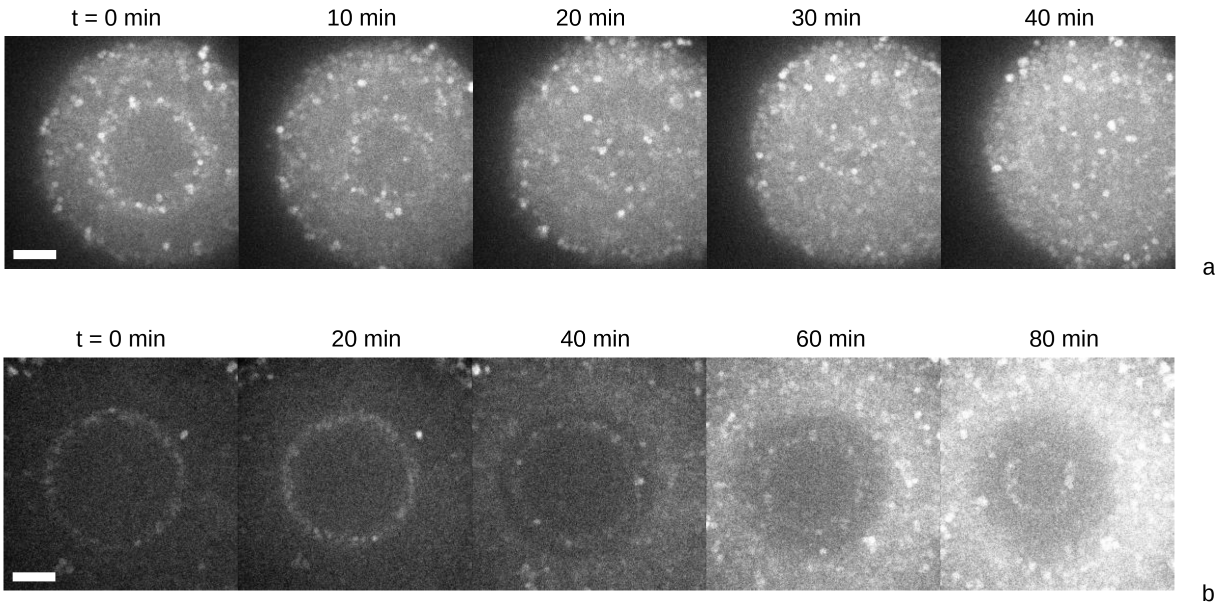


**Fig. S9 Dynamics of membrane potential after azithromycin treatment**. Strain $wt^*$ (NG150), flow chamber. Colonies were treated for 30 minutes with azithromycin at MICx100, $0.64\,\mu g/ml$ (a,b : two different colonies), and exhibit within 1 to 2 hours after treatment a collective hyperpolarization and shell reversal. Scale bar: 5 µm.

| Strain | Genotype | Source |
| --- | --- | --- |
| $wt^*$ (NG150) | *G4::aac* | (28) |
| $wt\ green$ (NG194) | *lctp:PpilE*<br>*sfgfp speR:aspC*<br>*G4::aac* | (32) |
| $\Delta pilT green$ (NG231) | *pilT::m-Tn3cm*<br>*lctp:PpilE sfgfp speR:aspC*<br>*G4::aac* | (27) |
| $pilT_{WB2}$ (NG176) | *iga::*$P_{pilE}$ $pilT_{WB}$ *ermC*<br>*G4::aac* | (25) |

**Table S1** Strains used in this study

### Movie captions

**Movie S1. Typical example of colony with blinking cell.** Movie corresponds to Fig. S2a. $\Delta t = 1$ min.

**Movie S2. Three-dimensional reconstruction of formation and propagation of the hyperpolarized shell.** $\Delta t = 5$ min.

**Movie S3. Typical example of onset of collective hyperpolarization for colonies with different initial sizes.** $\Delta t = 5$ min.
